# Differential photoperiodic control of morning and evening expressed transcripts in tomato

**DOI:** 10.64898/2026.03.06.710147

**Authors:** Alberto González-Delgado, Krzysztof Wabnik, José M Jiménez-Gómez

## Abstract

Photoperiod, the daily duration of light, is a key environmental cue that varies with season and latitude. Photoperiod signals profoundly influence plant growth and development, and play a central role in crop adaptation across latitudes. Deciphering how plants perceive these seasonal light cues at the molecular level is essential for understanding crop evolution and shaping their geographical distribution.

In this study, we investigated how photoperiod influences gene expression dynamics in tomato (*Solanum lycopersicum*) by performing an RNA-seq time-course across three photoperiod regimes, sampling at two-hour intervals. We use isogenic lines segregating for wild alleles of genes involved in circadian rhythm domestication in tomato to describe their contributions to the circadian clock and flowering time pathways.

The high temporal resolution of our experiment allowed precise characterization of transcriptional dynamics and their responses to photoperiod, including shifts in phase, amplitude, and waveform. We found that most transcripts time their expression to dawn or approximately 12 hours later, resulting in a systematic misalignment between evening transcripts and the actual timing of dusk. Morning- and evening-phased transcripts differ markedly in expression levels and waveform responses to changes in photoperiod. Together, these patterns suggest that morning transcripts sense photoperiod transcriptionally, and serve as zeitgeber references for evening transcripts, that could measure photoperiod length by coincidence with external cues. Together, our results provide new insight into the molecular basis of photoperiodic adaptation in plants.

## INTRODUCTION

The most reliable time cues (*zeitgebers*) on Earth arise from its rotation and orbit around the Sun, giving rise to daily light–dark cycles and seasonal changes. The duration of the light period within the 24-hour cycle, called photoperiod, varies predictably across seasons and latitudes. Most organisms have evolved mechanisms to synchronize their internal biological processes with these external cycles, generating endogenous ∼24-hour oscillations known as circadian rhythms. These rhythms confer adaptive advantages by aligning physiology and metabolism with environmental timing, thereby improving performance in fluctuating conditions. Photoperiodic responses are widespread across the plant lineage, and regulate key aspects of plant life cycles, including its time to flowering, growth, abiotic stress responses, defense against pathogens, and major aspects of plant metabolism, such as metabolic fluxes, photosynthate allocation, phytohormone homeostasis and photosynthesis efficiency (González-Delgado et al., 2025).

Photoperiod is perceived in plants through specialized photoreceptors that respond to light and temperature signals. These molecules undergo conformational changes upon stimulus and transmit signals through regulatory networks (Paik and Huq, 2019). Central among these networks is the circadian clock, a cell-autonomous transcriptional–translational feedback system generating self-sustained rhythms with an approximately 24-hour period (Brian Thomas and Daphne Vince-Prue, 1997). The clock integrates environmental inputs with internal metabolic feedback loops, including sugar signals (Frank et al., 2018; Haydon et al., 2013) and redox status (Stangherlin and Reddy, 2013), and controls both transcriptionally and post-transcriptionally the expression of hundreds of genes in a timely manner (Cai et al., 2025).

To align internal timekeeping with changing photoperiods, the plant circadian system undergoes continuous entrainment to light and temperature cycles along the duration of the year, primarily through tracking dawn, which is considered the *Zeitgeber*, the time-giver (Flis et al., 2016; Webb et al., 2019). Changing photoperiod exerts broad effects on the circadian clock and its downstream outputs, yet the nature and magnitude of these responses vary across species. As day length increases, *Arabidopsis thaliana*, potato, and quinoa show delayed transcript phases (timing of peak expression) and reduced oscillation amplitudes (difference between maximum and minimum expression) (Feke et al., 2025; Leung et al., 2023; Michael et al., 2008; Wu et al., 2023). In contrast, *Arabidopsis halleri* displays increased expression amplitudes under longer photoperiods (Nagano et al., 2019).

Photoperiod responses have profound agricultural relevance. Human selection for reduced photoperiod sensitivity has been crucial for expanding crop cultivation into new latitudes: nine of the ten crops with the largest harvested areas globally have undergone selection affecting their photoperiod responses during domestication or improvement (González-Delgado et al., 2025). In tomato, two key regulators of light integration into the circadian network, *EMPFINDLICHER IM DUNKELROTEN LICHT 1* (*EID1*) and *NIGHT LIGHT-INDUCIBLE AND CLOCK-REGULATED 2* (*LNK2*), were targeted during domestication. *EID1* encodes for an F-box E3 ligase known to interact with phytochromes (Marrocco et al., 2006) and circadian clock proteins such as ELF3 (Qin et al., 2023). Mutations in tomato’s *EID1* appeared early during domestication and delay circadian phase (Müller et al., 2018, 2016). *LNK2* encodes a transcriptional co-regulator that interacts with light-responsive REVEILLEs (RVEs) family of transcription factors in Arabidopsis. Through the formation of LNK-RVE8 complexes in the morning, LNK2 co-promotes the expression of evening-phased circadian clock genes, such as PSEUDORRESPONSE REGULATORS 5 and 3 (PRR5 and PRR3), TIMING OF CAB EXPRESSION 1 (TOC1) (Ma et al., 2018; Xie et al., 2014), GIGANTEA (GI), LUX ARRHYTMO (LUX) and EARLY FLOWERING 4 (ELF4). In contrast, LNK2-RVE8 complex is degraded in the evening, repressing morning-phased circadian clock genes including CIRCADIAN CLOCK ASSOCIATED 1 (CCA1), LATE ELONGATED HYPOCOTYL (LHY), and RVE8 (Hsu et al., 2013; Sorkin et al., 2023). In tomato, mutations in LNK2 delay the circadian period under constant conditions (Müller et al., 2018, 2016). Mutated alleles of EID1 and LNK2 likely allowed tomato’s expansion beyond its equatorial origins, making it an exceptional system for dissecting how photoperiod shapes transcriptomic rhythms.

In this work, we generate the most comprehensive temporal expression dataset in tomato to date across three photoperiods, including plants segregating for domestication-related alleles of EID1 and LNK2, and collecting samples at 2-hour intervals. We dissect global expression dynamics, study their principles and responses to photoperiod, and investigate how they are affected by the mutations driving tomato’s clinal adaptation.

## RESULTS

### Transcriptome-wide effects of diurnal cycles

Our experiment included four *S. lycopersicum* var *lycopersicum* isogenic lines: the cultivar MoneyMaker (MM; eid1/lnk2), and three lines carrying introgressions with wild *S. pimpinellifolium* alleles of *EID1* (line EID1, *EID1/lnk2*), *LNK2* (line LNK2, *eid1/LNK2*), and both *EID1* plus *LNK2* (line LE, *EID1/LNK2*) (Xiang et al., 2022). These lines were grown in one of three photoperiod regimes (Fig. 1A): short-day (SD: 6 h light/18 h dark), neutral-day (ND: 12 h light/12 h dark), and long-day (LD: 18 h light/6 h dark). All treatments were conducted at 21°C during the day and 18°C at night. Leaf samples were collected every two hours from two biological replicates per timepoint, genotype and photoperiod. On average, we obtained 32.7 million RNA-seq paired end reads per sample (range: 22.8–50.4 million), and after excluding low expressed transcripts, we analyzed expression values for 25,909 out of the 34,688 annotated transcripts (74.69%).

**Figure 1.**
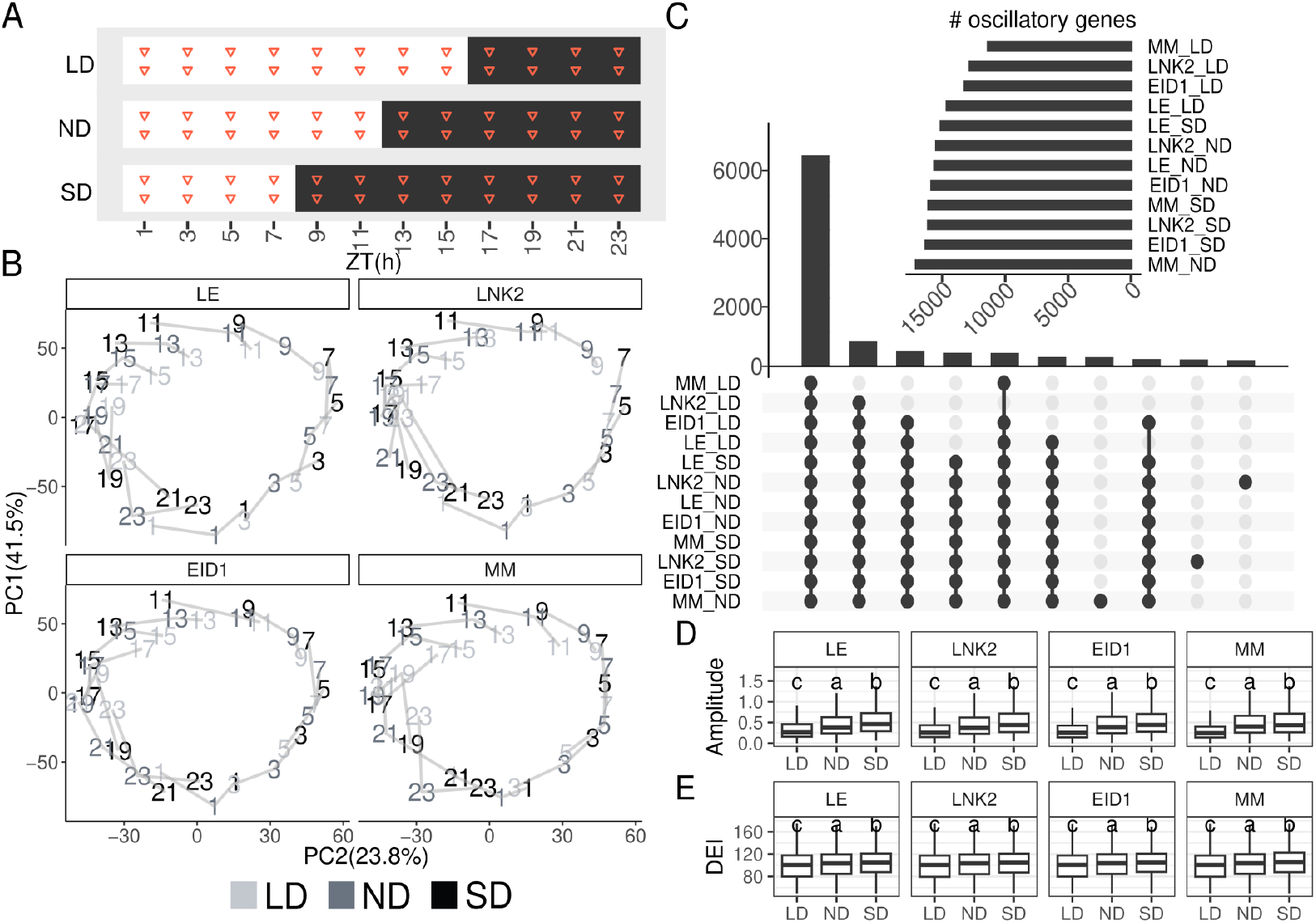
Detection of oscillations in the diurnal transcriptome. (**A**) Experiment timeline representation. Zeitgeber Times (ZT) for sample collection (red triangles) are indicated in the x axis. White and black boxes represent day and night, respectively. (**B**) Principal Component Analysis from average normalized expression values for each timepoint, genotype and condition. Numbers indicate timepoints (ZT), and colors represent photoperiods. Samples collected at similar ZT are linked with grey segments. (**C**) Top: Number of oscillating genes per condition and genotype. Bottom: Intersecting oscillating genes across condition and genotype. Transcript amplitude (**D**) and daily expression integral (E) values by photoperiod and genotype. Different letters above each box indicate significant differences between groups based on Tukey’s HSD test for each genotype separately (α = 0.05).

Principal Component Analysis (PCA) of transcriptome wide expression values revealed a circular trajectory with consecutive samples ordered counterclockwise (Fig 1B). This pattern confirms the consistency of our data and aligns with a transcriptomic wide diurnal progression that resets every 24 hours. It is noteworthy that time points from different photoperiods are systematically shifted relative to one another, with samples collected at a given ZT being significantly closer to samples taken two hours earlier in shorter photoperiods and two hours later in longer photoperiods (Fig1B; S1). This suggests that photoperiod primarily re-times genome-wide expression programs rather than disrupting their organization.

We detected oscillatory transcripts using autoregressive spectral analysis and harmonic regression models implemented in ARSER, which fit sinusoidal functions to the gene expression profiles (Wu et al., 2016). We found that 90.69% of expressed transcripts exhibited oscillatory behavior in at least one condition or genotype (FDR<0.01), underscoring that most biological processes are under strong temporal regulation (Fig 1C; Table S1). This high proportion of oscillating transcripts exceeds by 9.32% in MM and 14.29% in LE previous reports under comparable conditions using the cultivar M82 and the wild species *S. pennellii*, to which MM and LE are genetically equivalent, respectively (Müller et al., 2016) (Figure S2; Table S2). The increase in the number of oscillating genes in the current experiment is likely due to its higher temporal resolution (2 h vs. 4 h sampling).

Most cycling transcripts cycled in all photoperiods and genotypes (51.97% of expressed transcripts), but there was a reduced number of oscillating transcripts under LD conditions. This could arise from two non-exclusive mechanisms: (i) impaired circadian regulation of gene expression or (ii) a general dampening of transcript abundance, both of which would reduce the statistical power to detect rhythmicity. To explore these possibilities, we examined for each transcript both the amplitude of oscillation and the Daily Expression Integral (DEI), defined as the sum of expression values across the 24-hour cycle, restricting our analysis to transcripts that cycled in all three photoperiods to allow meaningful comparisons across treatments. These transcripts displayed markedly lower amplitudes and DEI values with longer photoperiods (Fig. 1D, 1F). Consistent with this trend, transcripts that oscillated in a greater number of conditions exhibited progressively higher amplitudes and DEI values (Fig. S3A, S3B). These results suggest that the underlying number of oscillating transcripts is likely similar in all photoperiods, but detection in LD fails for transcripts with lower expression levels and oscillation amplitudes.

### Temporal coordination of transcripts with light-dark cycles

We next examined the phase of oscillating transcripts and identified two prominent groups. Approximately half of rhythmic transcripts peaked near ZT0 ([ZT18–ZT6), called hereby MPTs for Morning-Phased Transcripts), while another half peaked near ZT12 ([ZT6–ZT18), called hereby EPTs for Evening-Phased Transcripts) (Fig. 2A).

**Figure 2.**
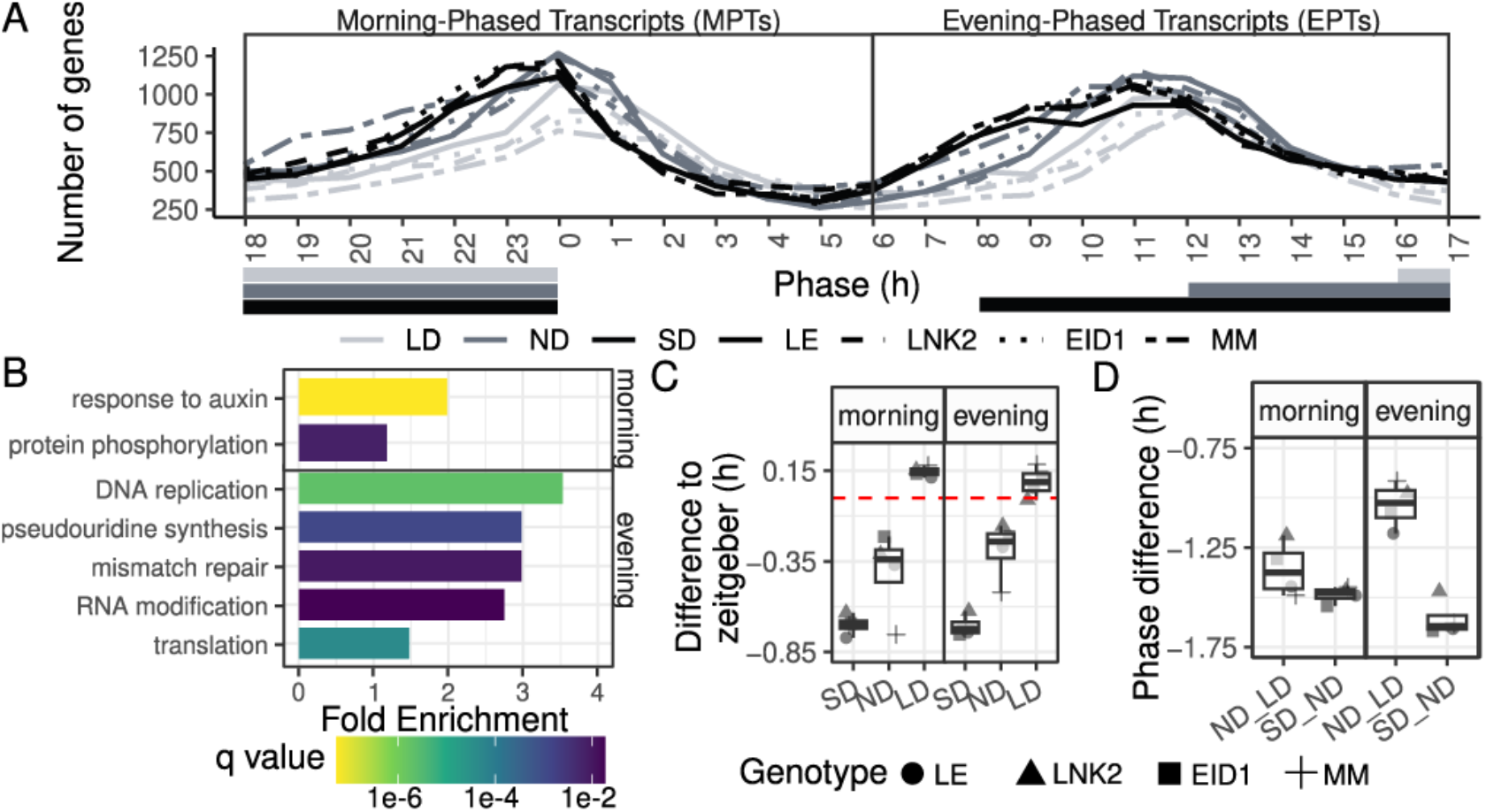
Photoperiod-dependent changes in the transcriptomic oscillation’s alignment of cycling genes. (**A**) Number of cycling genes (y axis) peaking at each hour of the day (x axis) across photoperiods and genotypes. Colored horizontal bars below the x axis indicate night segments. (**B**) Gene Ontology (GO) terms significantly enriched in MPTs (top) and EPTs (bottom). Fold enrichment values and adjusted q-value in a hypergeometric test are showed in the x axis and fill color respectively. (**C**) Distance between each transcript’s phase and its closest Zeitgeber time (red dashed line): being ZT0 for MPTs (left panel) and ZT12 for EPTs (right panel). (**D**) Short-day and neutral-day (SD–ND), or long-day and neutral-day (LD–ND) phase differences for MPTs (left panel) and EPTs (right panel).

Functional enrichment analysis revealed clear temporal partitioning of biological processes between groups. MPTs were enriched for processes such as response to auxin and protein phosphorylation (Fig. 2B). In contrast, EPTs were enriched for translation, RNA modification, mismatch repair, pseudouridine synthesis and DNA replication (Fig. 2B). This indicates coordination of growth-related processes in the morning and genome maintenance processes later during the day. Motif enrichment analyses showed the canonical Evening Element (EE) strongly enriched among EPTs, consistent with direct regulation by the evening complex (Table S3). The morning element motif was also enriched among MPTs, although not prominently (Table S4).

Circadian parameters of EPTs and MPTs were mostly similar (Fig. S4). In both groups, the closer a transcript peaks to ZT0 or ZT12, the higher the amplitude of their oscillations, the closer their periods to 24 h, and the higher the adjusted R-square in the harmonic regression model compared to transcripts peaking further away from these times. This suggests stronger circadian control on transcripts peaking at ZT0 and ZT12. The only visible difference between EPTs and MPTs was their overall expression level, with higher DEI values in the morning (Fig. S4D), suggesting stronger regulatory activity during the preceding night.

We then studied how photoperiod affects phase for individual transcripts. Most transcripts timed their maximum expression to occur between 0.01 and 0.77 hours of ZT0 or ZT12 (Fig. 2C). They experienced an average delay of 1.37h as photoperiod increases by 4 hours, with variation well below one hour between photoperiods and groups (Fig. 2D). This indicates that the 4-hour changes in the light regime do not shift the phase of expression proportionally, and creates a misalignment of EPTs with their evening *Zeitgeber* in LD and SD: While MPTs remain tightly aligned with dawn across all photoperiods, EPTs align with dusk only under ND conditions, and peak their expression during dark periods in SD and during light periods in LD (Fig. 2A). This misalignment positions ETPs, and possibly their encoded proteins, to respond to light availability through external-coincidence mechanisms.

### Photoperiod-dependent modulation of transcriptional dynamics

To study the expression dynamics of EPTs and MPTs, we selected high amplitude transcripts peaking within 30 minutes of ZT0 and ZT12, and called them ZT0s and ZT12s respectively (Fig 3). ZT0s displayed higher expression profiles than ZT12s, and their expression profiles also seemed more affected by the photoperiod (Fig 3A). We used Functional Data Analysis (FDA) to study these waveform differences and computed their first (d1) and second (d2) derivative components, which capture the rate of expression change and the curvature (or acceleration) of those changes, respectively. In ZT0s, the d1 component showed strong photoperiod-dependent variation, particularly throughout the rising phase that coincides with the dark (Fig. 3B). ZT12s displayed much smaller differences, that also coincide with the dark, but corresponds to their inhibition phase. These patterns suggest that the integration of day-length information occurs primarily during the dark period.

**Figure 3.**
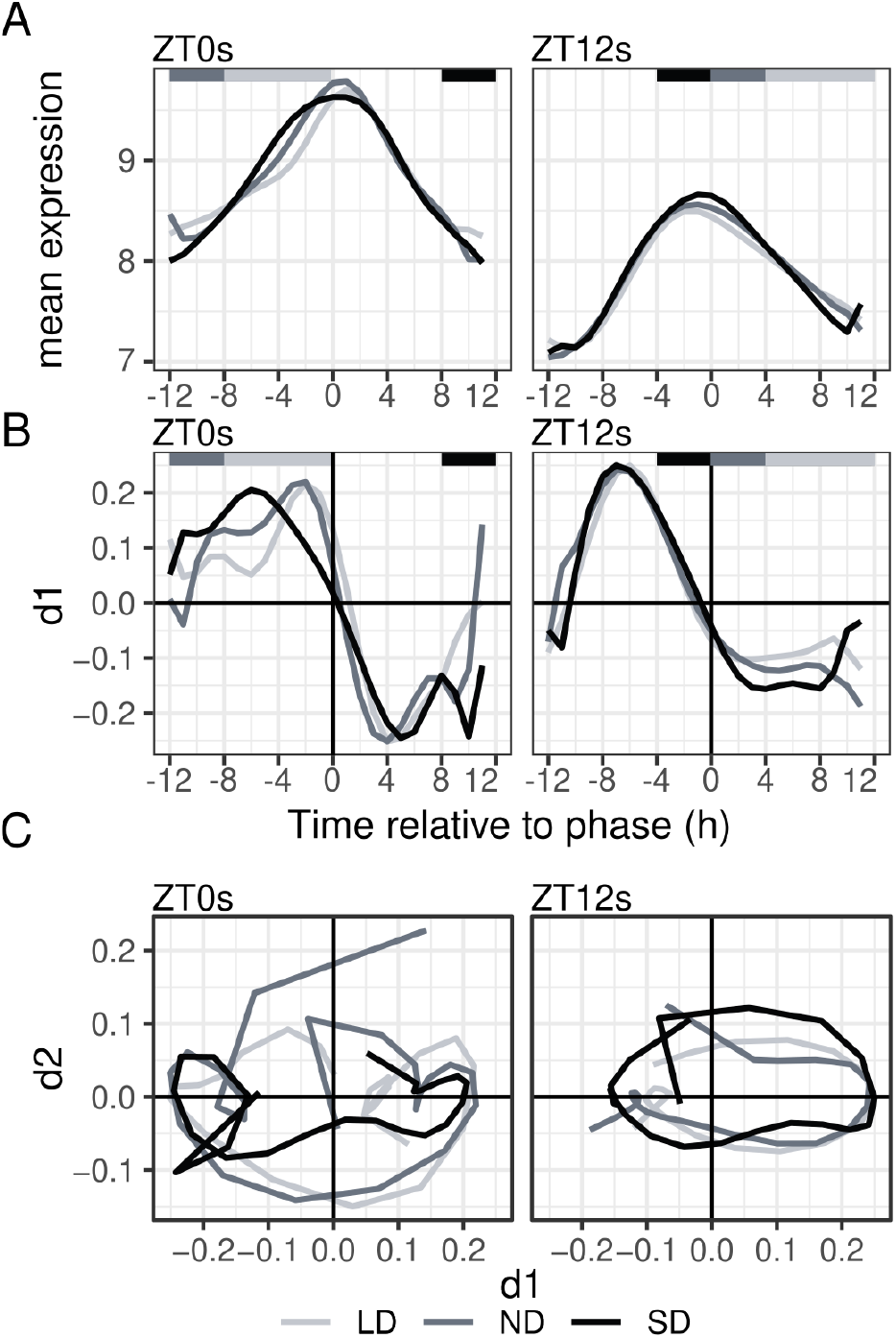
Photoperiodic temporal dynamics of MPTs and EPTs. Columns in each panel represent average values per photoperiod for transcripts with phases within 30 minutes of ZT0 (ZT0s, left) and ZT12 (ZT12s, right). For all calculations, variance-stabilized expression values were centered to the predicted transcript phase (0 in all x axes). (**A**) Average expression per photoperiod (**B**) First temporal derivative (d1) from FDA analysis. Left of the peak corresponds to the rising phase and right to the falling phase. (**C**) Trajectories for the first (d1, x-axis) and second (d2, y-axis) derivatives.

We then considered trajectories in d1–d2 space (Fig 3C). ZT12s, showed constrained, round-shaped loops in all photoperiods, reflecting constant and stable oscillatory dynamics. In contrast, ZT0s presented open circles with two small, symmetrically opposed loops, reflecting two distinct oscillatory modes, one favored in LD and the other in SD (positive and negative values of d1, respectively). These photoperiod-dependent changes in trajectory geometry alter the time of maximum expression of ZT0s, advancing their phase in SD and delaying it in LD, in comparison with the more stable oscillatory dynamics in ZT12s. All results from ZT0s and ZT12s were confirmed in a broader set of MPTs and EPTs (Figure S5).

In summary, we find that MTPs, despite being expressed under similar environmental conditions in all photoperiods, change their waveforms in response to photoperiod, while ETPs are significantly less affected, and are expressed in radically different environmental conditions in SD (dark), ND (zeitgeber), and LD (light). This suggests a model with distinct regulatory properties of MTPs and ETPs, with MTPs responding transcriptionally to photoperiod, potentially using timing information provided by the coincidence of ETPs with environmental signals.

### Differential roles of EID1 and LNK2 in photoperiodic responses

We had observed in Figure 1C that LD reduced the number of oscillating transcripts. This reduction depends on the genotype in a pattern that might be relevant for domestication. MM presented fewer oscillating genes in LD than LE, with the EID1 and LNK2 lines being intermediate (Fig. 1C). These differences were shared both by EPTs and MPTs (Fig S6A). In order to study the regulatory pathway leading to these differences, we analyzed the promoter of genes that oscillate in LE but not in MM in MPTs and EPTs (FigS6B). We found a significant enrichment of the REF6 binding motif specifically among the MPTs (FigS6B). REF6 is known to physically interact with PHYB to regulate gene expression (Yan et al., 2025). Given that EID1 genetically interacts with PHYB1 in tomato (Müller et al., 2018), we hypothesize that loss of EID1 function may lead to altered PHYB availability, thereby contributing to the observed differences in diurnal gene expression. Notably, this motif is also the most enriched in the MPTs oscillating in EID1 but not in LNK2, reinforcing the link between EID1 and REF6 regulating gene expression in LD (FigS6C, S6D).

We then assessed how LNK2 and EID1 influence the expression of individual genes. For LNK2, we identified 1,243 differentially expressed genes (DEGs) at one or more timepoints throughout the day between plants carrying the cultivated and wild alleles. The majority of DEGs peaked around ZT11 (Figure S7), which is the timing of LNK2 activity in Arabidopsis (Sorkin et al., 2023). Upregulated transcripts were enriched in cell redox homeostasis, microtubule-based movement and DNA replication GO terms (Figure S8). Downregulated transcripts were enriched in regulation of DNA-templated transcription, transmembrane transport, cell redox homeostasis, photosynthesis, microtubule-based movement and photosynthetic electron transport in photosystem II. In *Arabidopsis thaliana*, the LNK1 protein binds to RVE8 and regulates the expression of circadian clock genes (Hsu et al., 2013; Sorkin et al., 2023).We found 16 circadian clock DEGs in LD, 15 in ND and 12 in SD. Among these, targets of the RVE8-LNK family complex in Arabidopsis such as PRR3, PRR5, PRR7, PRR9, RVE4/8, GI, CCA/LHY, ELF4, and TOC1 (Figure S9A). Notably, LNK2 deficiency alters the expression of core clock transcripts depending on their phase, reducing the expression of EPTs such as PRR5, GI, and TOC1, but increasing the expression of MPTs such as RVE4/8, PRR9 and CCA/LHY.

EID1 is a negative regulator of the PHYA signaling pathway in *Arabidopsis thaliana* (Büche et al., 2000) and controls flowering time in Soybean (Qin et al., 2023). We found 928 DEGs between plants with contrasting EID1 alleles at one or more timepoints (Figure S7). Upregulated genes were functionally enriched in translation, lipid metabolism, cell redox homeostasis, cell wall macromolecule catabolic process, chitin catabolic process, photosynthetic electron transport in photosystem II, DNA replication and response to wounding (Figure S8). Downregulated transcripts were enriched in transmembrane transport, lipid metabolism, response to wounding, photosynthetic electron transport in photosystem II, microtubule-based movement, intracellular signal transduction and sucrose metabolism. Two CENTRORADIALIS/TERMINAL FLOWER 1/SELF-PRUNING (CETS) flowering time regulators were present among DEGs (Fig S9B). These transcripts have an important role governing the life history of a plant, and are highly regulated by photoperiodic queues (Cao et al., 2016). Among these, the flowering time repressor SP5G was generally downregulated in ND when plants carry active EID1 alleles, which could not be observed in any other photoperiod. This gene is an example of the complexity of photoperiod regulation of expression dynamics in tomato. It gradually increases its expression with longer photoperiods, and shows a bi-phasic expression pattern with phenotypic relevance in other plants (Lee et al., 2023). SP11B.1 is a positive regulator of flowering from the CETS family, also shows photoperiod dependent expression, and it is upregulated in ND when EID1 alleles are active. The effect of EID1 increasing the expression of an activator of flowering (SP11B.1) and decreasing the expression of a negative regulator (SP5G) suggests a role of EID1 accelerating flowering time under ND, possible through its putative role degrading PHYB1.

Together, these results suggest that, in our conditions, deleterious mutations in LNK2 have a broader effect than in EID1, and impact especially the circadian clock, while mutations in EID1 have a more specialized function as phytochrome regulators and flowering time activator.

## DISCUSSION

### Role of LNK2 and EID1 in photoperiodic circadian regulation

Here, we studied variation in diurnal expression in tomato having into account its domestication history by using lines segregating in its two main circadian regulators, EID1 and LNK2. Previous work reported global differences in phase of expression between cultivated tomato and the wild species *S. pennellii*, which carries ancestral alleles of EID1 and LNK2, similar to LE (Müller et al., 2016). Expression phase shifts associated with the EID1 mutation were also predicted using the molecular timetable method from a single ZT2 sample in the same lines examined here (Xiang et al., 2022). However, we did not detect a global phase shift linked to either EID1 or LNK2 in our dataset. A plausible explanation for this is the difference in growth temperature between studies: 24 °C previously versus 21 °C in our experiment. Supporting this possibility, the effect of EID1 is temperature-dependent in Arabidopsis (Tu et al., 2024), and its function in tomato requires light and the photoreceptor PHYB1, which is also implicated in temperature responses (Müller et al., 2016). It is therefore possible that the lower temperature used here dampened the global impact of EID1. Nevertheless, both EID1 and LNK2 retained detectable effects on their known genetic targets: LNK2 influenced circadian clock gene expression, while EID1 modulated photoperiod-dependent florigen transcripts, as it has been described in Soybean (Qin et al., 2023). These findings suggest that, while EID1 and LNK2 can influence specific downstream pathways, their global impact on transcriptome phase is conditional by environmental context.

### The global transcriptome is modulated by photoperiod

The circadian system operates through a plastic network that continuously adjusts gene expression to optimize metabolic and developmental outputs depending on external cues (Webb et al., 2019). Our results demonstrate that photoperiod length strongly modulates the global phase of the transcriptome, advancing or delaying the average expression peak under shorter or longer days, respectively. The systematic phase shifts observed in the PCA indicate that, at the global scale, diurnal expression programs are preserved across photoperiods but occur earlier or later depending on day length. This was in contrast with the low number of cycling transcripts detected in LD compared with ND or SD, that suggests a disruption of circadian regulation under those conditions. However, our analyses of cycling parameters revealed significant reductions in amplitude and DEI in LD conditions, leading us to conclude that the low number of oscillating transcripts detected in LD likely reflects reduced sensitivity to weaker rhythms rather than a breakdown of the underlying circadian program. This result suggests that a comparable number of genes remain under circadian control across all photoperiods, but that rhythmicity becomes increasingly attenuated with extended light in tomato. Lower amplitude in longer photoperiods has been found in other species, like *Arabidopsis thaliana* (Leung et al., 2023), potato (Feke et al., 2025) or Quinoa (Wu et al., 2023) but the opposite is also true in *Arabidopsis hallerii* collected in natural conditions (Nagano et al., 2019) and *Ostreococcus tauri* (Romero-Losada et al., 2025). For this reason, the attenuation of transcript oscillation observed in LD could be dependent on the species and/or experimental conditions.

### Crosstalk between morning and evening genes integrates photoperiod

The 2-hour resolution in our dataset allows us to estimate circadian parameters with precision. We observed a robust bimodal distribution of the phase of cycling transcripts, with approximately equal amounts of transcripts peaking in expression close to ZT0 and ZT12. This pattern of expression has been observed frequently but not in all plant lineages (Ferrari et al., 2019). As more studies incorporate high-frequency sampling across diverse environmental conditions, it will become clearer how widespread these patterns truly are.

The bimodal distribution of phases allows partition of biological processes into two distinct temporal programs, with MPTs participating in growth and energy-demanding cellular activities, while EPTs working mainly in genome maintenance. The vast majority of MPTs and ETPs showed fairly similar circadian characteristics, and responded very similarly to photoperiod, indicating the effect of an internal circadian clock that times their expression to ZT0 and ZT12. In the case of EPTs, this means a misalignment with the time of lights off that allows environmental regulation through light or temperature coincidence mechanisms.

The major differences between MPTs and EPTs were characterized by using phase-plane qualitative analysis (Figure 3C). The global behavior of EPTs showed a closed orbit, indicative of a limit cycle with self-sustaining oscillations that are robust to disturbances. In this case, photoperiod would introduce modulation without changing the dynamic regime, resulting in quantitative changes in transcript amplitude. In contrast, MPTs show small loops in the phase plane, corresponding to spiral equilibrium points typical of dampened or controlled systems. In these systems, the trajectory does not return to the same state, reflecting dissipation or external forcing, and follows transient oscillatory paths characteristic of controlled systems with alterations in local stability and regulatory couplings. These changes in the attractor landscape cause qualitative modifications in the loops’ structure. Together, these observations highlight a clear distinction between the robust limit cycles of EPTs and the more flexible, photoperiod-sensitive dynamics of MPTs.

This stronger photoperiod-dependent waveform changes in MPTs allow us to speculate on how photoperiodic information is integrated in tomato. Particularly, the waveform alterations observed in MPTs occurred during gene induction, where they can impact the time at which a transcript reaches maximum expression (their phase). However, the waveform alterations in EPTs were minor, and they occur during its inhibition phase, which does not affect the time of maximum transcript abundance. This pattern suggests a model in which MPTs integrate day-length information by changing their activation rate and subsequently their phase, and act as a temporal reference for EPTs that would merely peak approximately 12 hours later (Figure 4). However, the photoperiod responses in waveform in MTPs cannot be direct result of different environmental cues, since all these transcripts peak very close to dawn independently of photoperiod. Instead, it could be elicited by ETPs and their encoded proteins, whose photoperiod-dependent coincidence with light determines their effect on MPTs’ expression. This is reminiscent of the functioning of some ETPs in Arabidopsis, that are central to the circadian clock and the photoperiodic regulation of flowering time, like CONSTANS, GI and ELF3, whose light dependent activity in the afternoon determine flowering time, and controls the expression of downstream circadian clock MPTs, like CCA1 and LHY1 (Anwer et al., 2020; de los Reyes et al., 2024). In summary, our results support a model in which MPTs act as primary integrators of photoperiodic cues at the transcriptional level, while this environmental information is perceived and transmitted at the post-transcriptional level by EPTs.

**Figure 4.**
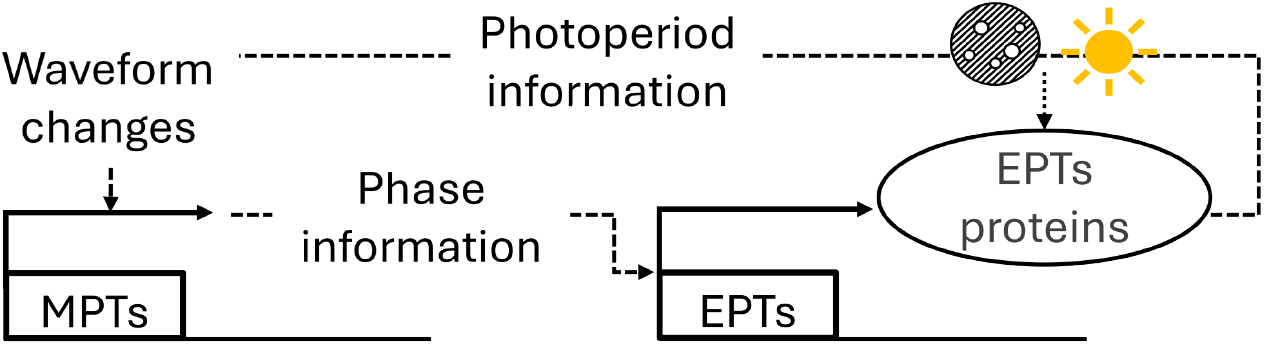
Proposed model for photoperiod integration and phase shifting. Morning-phased transcripts (MPTs) adjust their expression waveform in a photoperiod-dependent manner, which changes their phase and that of the evening-phased transcripts (EPTs), through regulatory cascades. EPT-encoded proteins integrate day-length information via external coincidence mechanisms, and this integrated signal is conveyed back to morning transcripts, resulting in photoperiod-dependent remodeling of their waveform.

## MATERIALS AND METHODS

### Plant material and RNA-Seq analysis

Tomato plants were grown under three different photoperiod conditions, including SD (short days: 6 hours light/18 hours dark), LD (long days: 18 hours light/6 hours dark), and ND (neutral days: 12 hours light/12 hours dark). In all cases, seeds were geminated in water in petri dishes in the dark, planted in soil after germination, and grown for one week in LD conditions. After this, they were moved to their respective photoperiod, where they stayed for 2 weeks. Leave samples were collected every two hours starting one hour after dawn (ZT1) and finishing 22 hours later (ZT23, Figure 1A). Total RNA was extracted using the RNeasy Plant Mini Kit from Qiagen. Sequencing libraries were prepared according to the Illumina TruSeq RNA protocol and sequenced in two lanes of a Novaseq 6000 system. Reads for each biological replicate were independently aligned to the tomato genome reference sequence v4.1 using hisat2 with default parameters (Kim et al., 2019). The number of reads per transcript in the ITAG4.1 annotation (Fernandez-Pozo et al., 2015) was obtained with the featureCounts function in the Rsubread R package (Liao et al., 2019). An average of 90.6% of the reads (range 84.2% to 92.8%) in each sample aligned uniquely to the reference. Normalized expression values per transcript were obtained by variance stabilizing transformation in DESeq2 (Love et al., 2014).

### Description of Circadian rhythms

Transcripts exhibiting circadian rhythms were identified on vst normalized expression values using ARSER, available in the Metacycle R package (Wu et al., 2016). ARSER requires a single expression value per gene and performs best with time-course data extending beyond 24 hours. To meet these requirements, we averaged our two biological replicates per timepoint, genotype and condition, and we duplicated the resulting data matrix to approximate a second day of measurements, assuming equivalency of two consecutive days in our controlled-environment chambers.

Oscillating genes were identified using the Benjamini–Hochberg adjusted p-value (FDR < 0.01). For each gene, we retained its period, phase, amplitude, and the cosine-fit quality (Adjusted R2) and coefficient of variability estimated by ARSER. We additionally computed DEI, defined as the sum of expression values across the 24-hour cycle.

To capture gene expression dynamics, genes that oscillate in all three photoperiods in each genotype were classified as morning-phased transcripts (MPTs) with phases in the interval [18,6) or evening-phased transcripts (EPTs) with phases in the interval [6,18). For expression dynamic studies, the time-course was centered on their phase estimated by ARSER, with all genes peaking exactly at 12. Variance-stabilized expression values were smoothed a fine temporal grid with 0.1 h intervals, and their first and second derivatives calculated FDA with B-spline basis functions included in fda R package (Ramsay et al., 2025).

### Differential expression analysis

Differential Expression analysis was performed with raw read counts using the DESeq2 package (Love et al., 2014). Normalization of the data was carried out using DESeq2 default method, which adjusts for differences in sequencing depth by calculating size factors for each sample. Samples were compared by their genotype at EID1 or LNK2 at each timepoint and condition using their cultivated alleles as a reference. The Benjamini-Hochberg method was used to control false discovery rate (FDR). Absolute values of Fold-Change higher than 1 and FDR lower than 0.05 were considered differentially expressed. The identifiers for circadian clock genes and CETs used in this analysis are reported in Table S5.

### Motif Enrichment Analysis

Motif Enrichment Analysis was performed with the script findMotifs.pl in Hypergeometric Optimization of Motif EnRichment (HOMER) tool (Heinz et al., 2010) using the genome and annotation of *Solanum lycopersicum* (version ITAG4.1). Motifs were searched within 1500 bp upstream of their transcription start sites

## DATA AVAILABILITY

The RNA-seq raw data underlying this article are available in Sequence Read Archive (SRA) at https://www.ncbi.nlm.nih.gov/sra, and can be accessed with the unique identifier PRJNA1180479.

## ACKNOWLEDGEMENTS

We thank the Adaptive Genetics and Genomics group and the Synthetic Biology of Plant Signaling Circuits group at CBGP for their helpful discussions, and their intellectual, technical and moral support. We also thank the CBGP and CEPLAS (Cluster of Excellence on Plant Sciences) for creating a program to finance AGD, and specifically Björn Usadel and Shizue Matsubara for their help making possible this project, and to and Marie Bolger for hosting AGD in their facilities at Julich Forschungszentrum. We thank the Plant Observatory Facilities at IJPB, INRAE, Versailles, for taking care of the controlled-environment growth facilities used in this work, and the Saclay Plant Sciences Network for enabling sequencing of all samples.

## AUTHOR CONTRIBUTIONS

A.G.-D. performed data analyses, generated figures, and wrote the initial draft of the manuscript. K.W. contributed to supervision, provided guidance on computational analyses, and assisted with manuscript preparation. J.M.J.-G. generated the RNA-seq data, provided scientific supervision, contributed to manuscript writing, and supported data interpretation and analysis. All authors reviewed and approved the final manuscript.

