## Supplementary Figures for "Differential photoperiodic control of morning and evening expressed transcripts in tomato"

### Figure S1

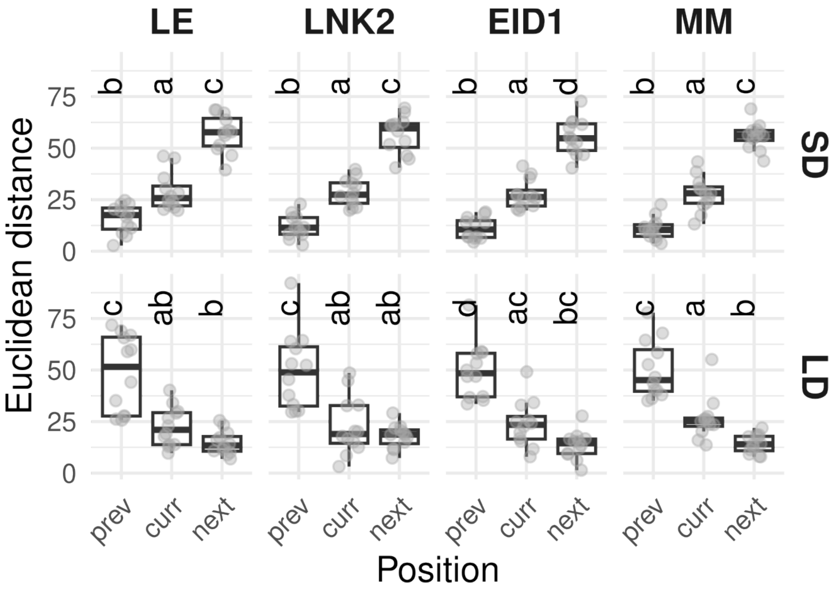


**Figure S1: Euclidean distances in PCA space between timepoints in different photoperiodic conditions.** The Euclidean distance between PCA values in ND and SD (top) or LD (bottom) at three relative Zeitgeber time (ZT) points: the previous (prev), current (curr), and next (next) time point. Different letters above boxes indicate significant differences among groups based on Tukey's HSD test (α = 0.05) per genotype. The graph shows that PCA values in ND are closer to previous timepoints in SD and later timepoints in LD.

**Figure S2**

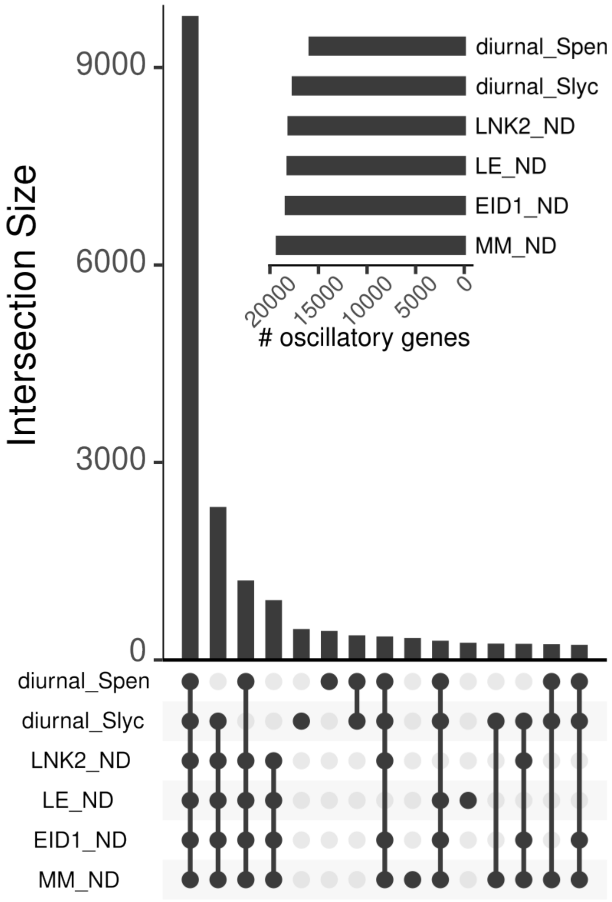


**Figure S2: Number of cycling transcripts detected in our current experiment versus Müller et al., 2016.** Previous dataset is labeled as diurnal Spen (*Solanum pennellii*) and Slyc (*Solanum lycopersicum*), and new dataset is labeled as genotype_photoperiod. Top: Number of oscillating transcripts. Bottom: Intersection of oscillating transcripts, highlighting shared and unique rhythmic transcripts.

### Figure S3

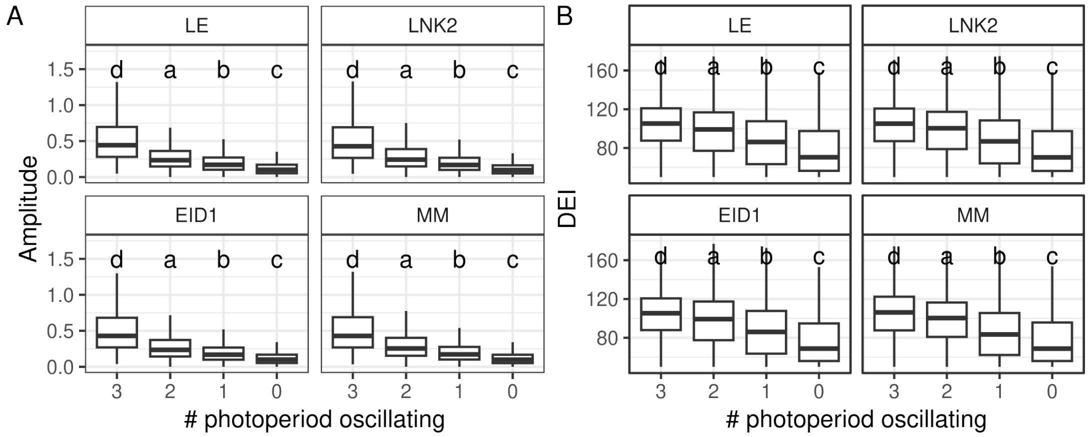


**Figure S3: Transcript amplitude (A) and Daily Expression Integral (DEI) (B).** Genes are grouped by the number of photoperiods in which they oscillate (3, 2, 1, or 0). Letters above boxes indicate significant differences between groups based on Tukey's HSD test (α = 0.05) in each genotype separately.

**Figure S4**

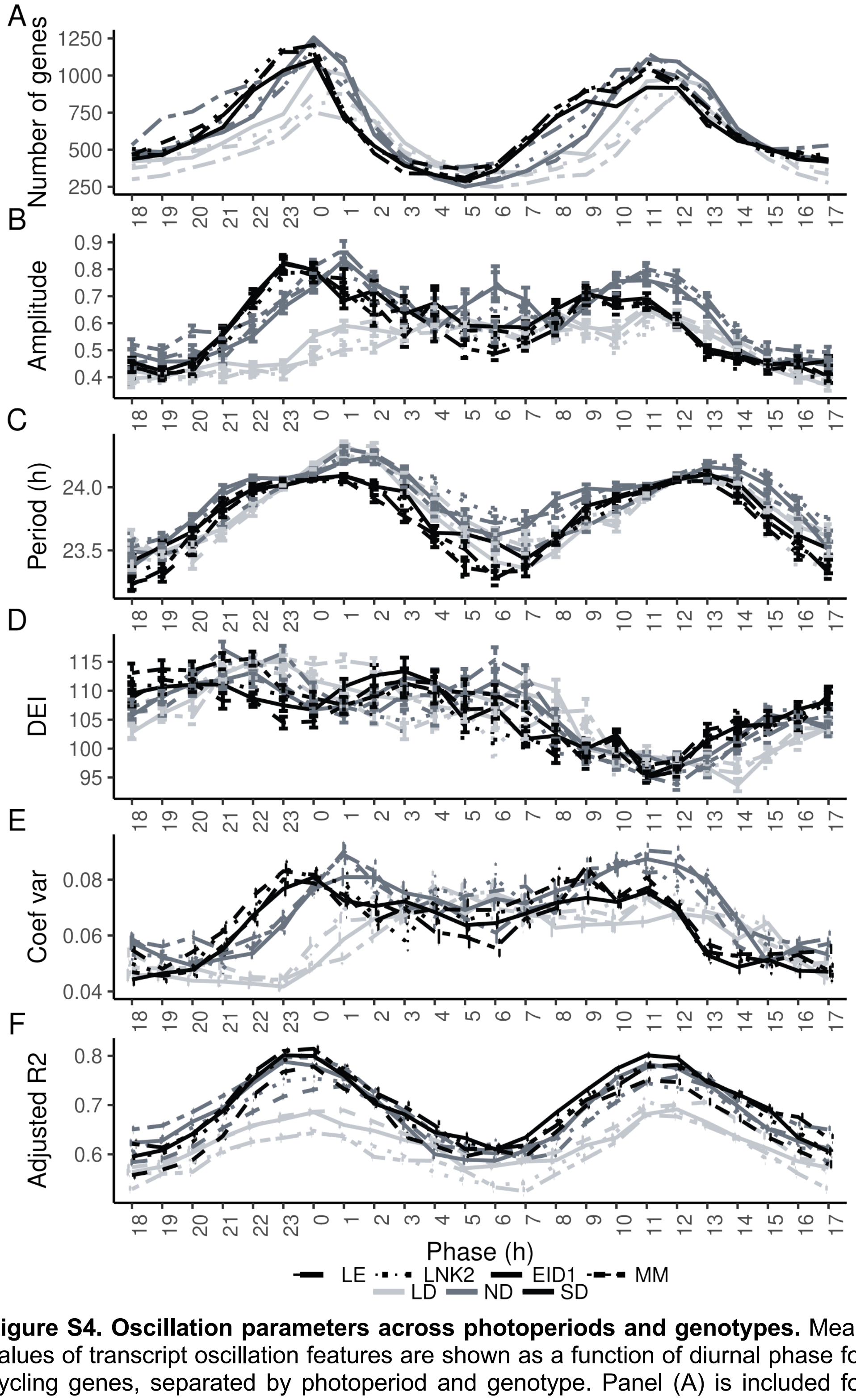


**Figure S4. Oscillation parameters across photoperiods and genotypes.** Mean values of transcript oscillation features are shown as a function of diurnal phase for cycling genes, separated by photoperiod and genotype. Panel (A) is included for guidance and represents the number of cycling genes (y axis) with phase at each ZT (x axis). For panels B, C, D, E and F, the y axis represents the average descriptor value for genes peaking at each phase bin (x axis).

**Figure S5**

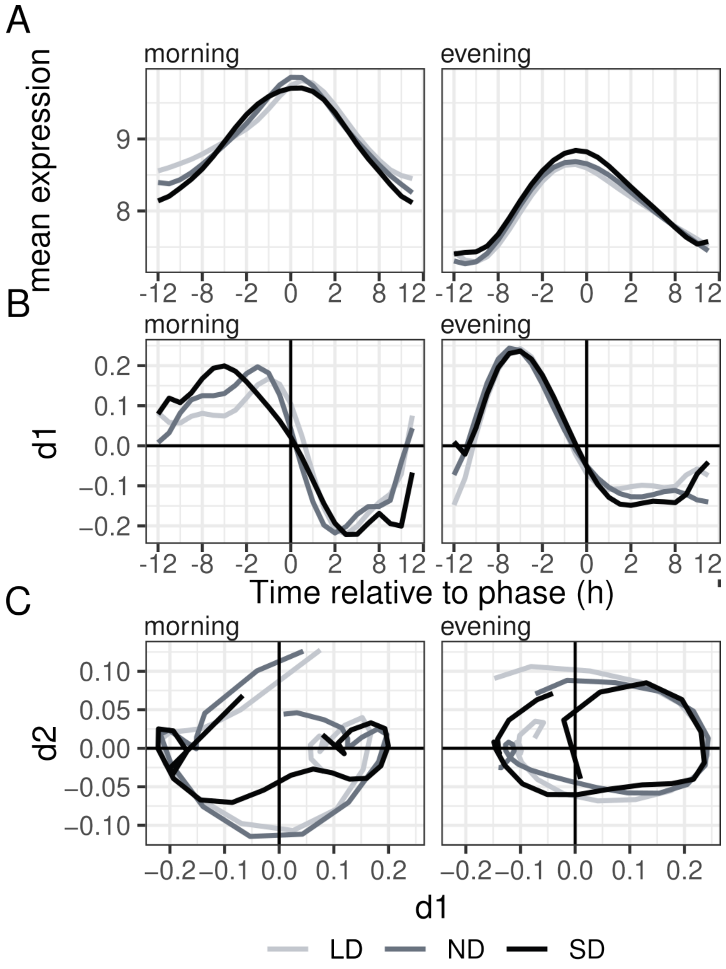


**Figure S5. Photoperiodic temporal dynamics of MPTs and EPTs.** Columns in each panel represent average values per photoperiod for transcripts with phases within 2 hours and 30 minutes of ZT0 (morning, left) and ZT12 (evening, right). For all calculations, normalized expression values were centered to the predicted transcript phase (0 in all x axes). (**A**) Average expression per photoperiod (**B**) First temporal derivative (d1) from FDA analysis. Left of the peak corresponds to the rising phase and right to the falling phase **(C)** Trajectories for the first (d1, x-axis) and second (d2, y-axis) derivatives.

**Figure S6**

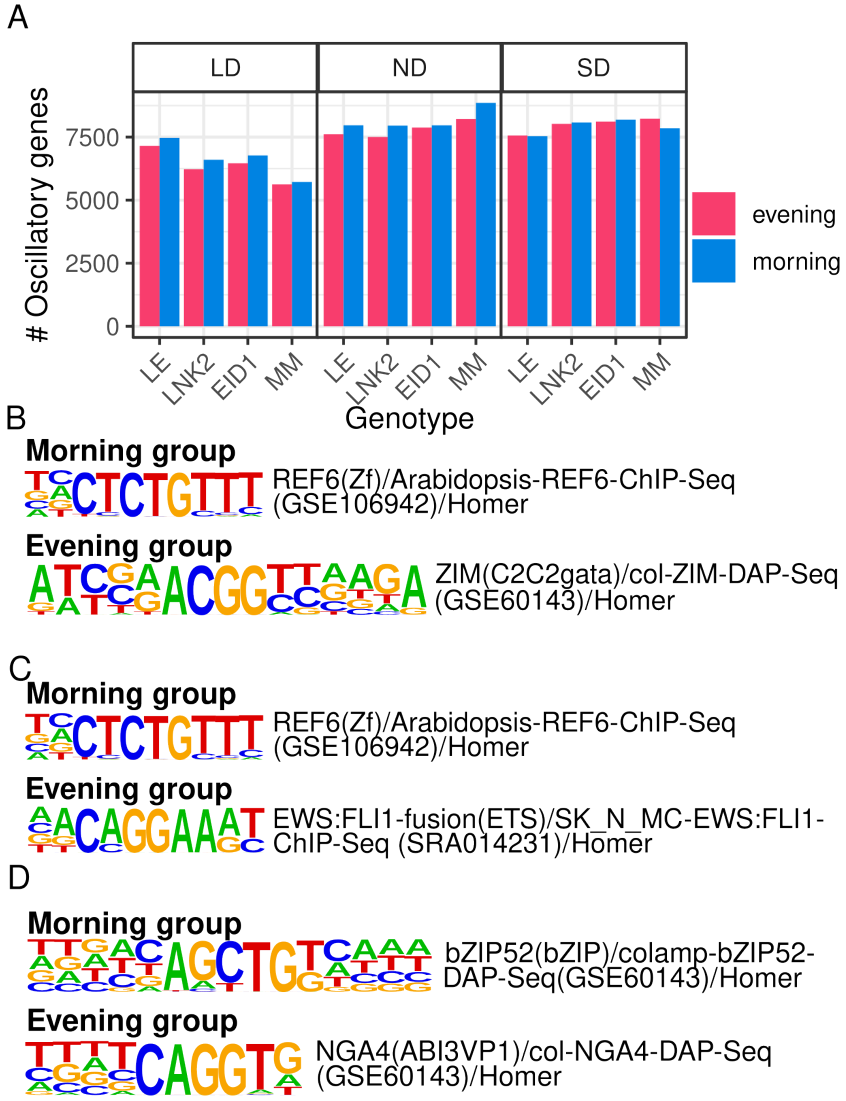


**Figure S6: LD-specific effect for EID1 and LNK2 on the number of cycling genes.** (A) Number of oscillatory transcripts by genotype, condition and phase group. Sequence logos show significantly enriched motifs in promoters of morning and evening transcripts under LD conditions oscillating in LE but not in MM (B), oscillating in EID1 but not in LNK2 (C) or oscillating in LNK2 but not in MM (D).

**Figure S7**

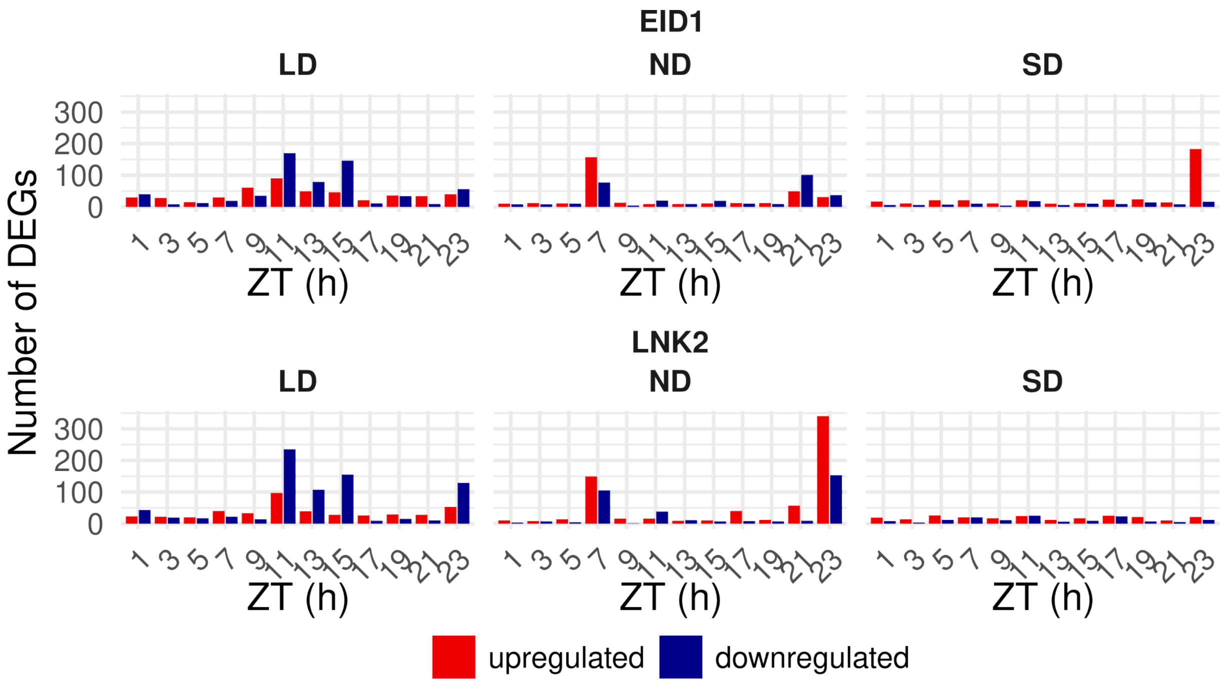


**Figure S7: Number of differentially expressed transcripts per time point by EID1 and LNK2.** Number of differentially expressed genes between genotypes grouped by their EID1 (top) or LNK2 (bottom) alleles at each Zeitgeber Time (x-axis). Upregulated genes are depicted in red and downregulated genes in blue.

### Figure S8

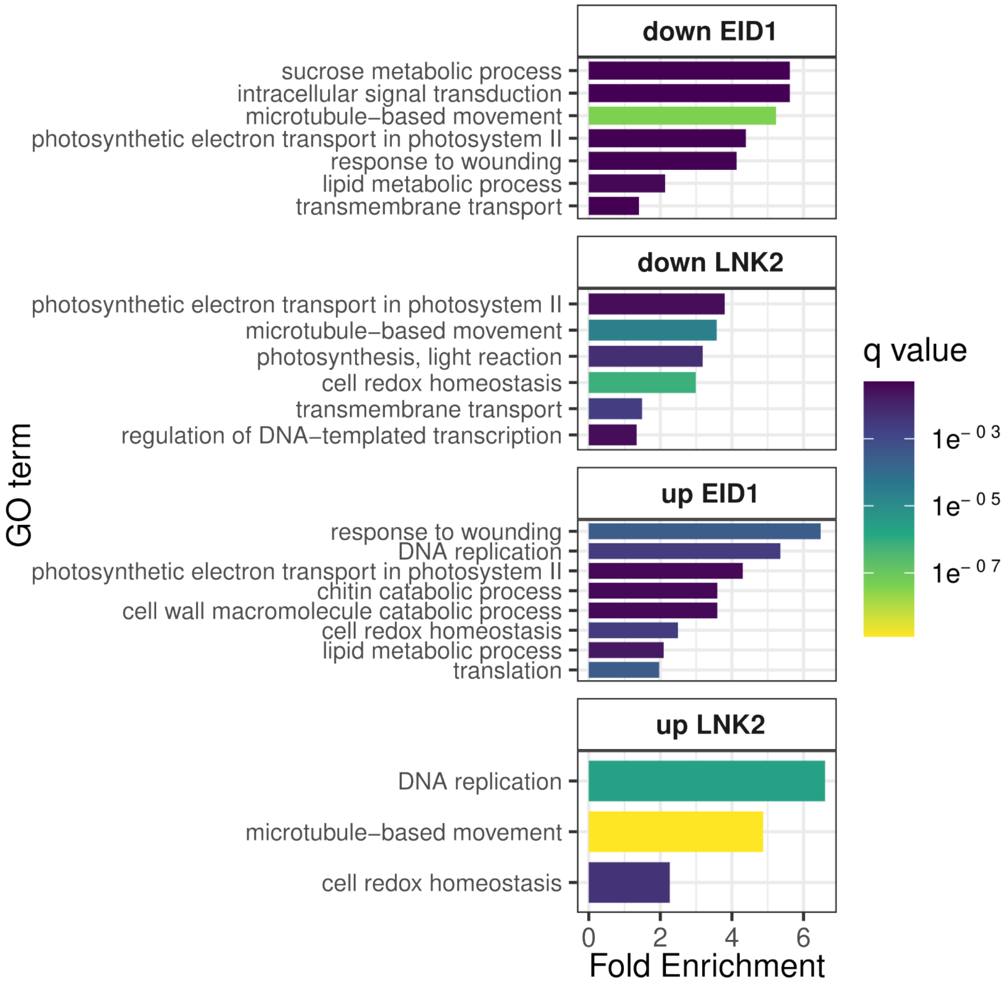


**Figure S8. Gene Ontology enrichment of differentially expressed genes by LNK2 and EID1.**
Gene Ontology (GO) enrichment analysis of differentially expressed genes (DEGs) identified in comparisons between genotypes grouped by their LNK2 (bottom) and EID1 (top), separated into downregulated and upregulated sets. Fold enrichment and the adjusted q-value from a hypergeometric test are indicated in the x-axis and fill color, respectively.

**Figure S9**

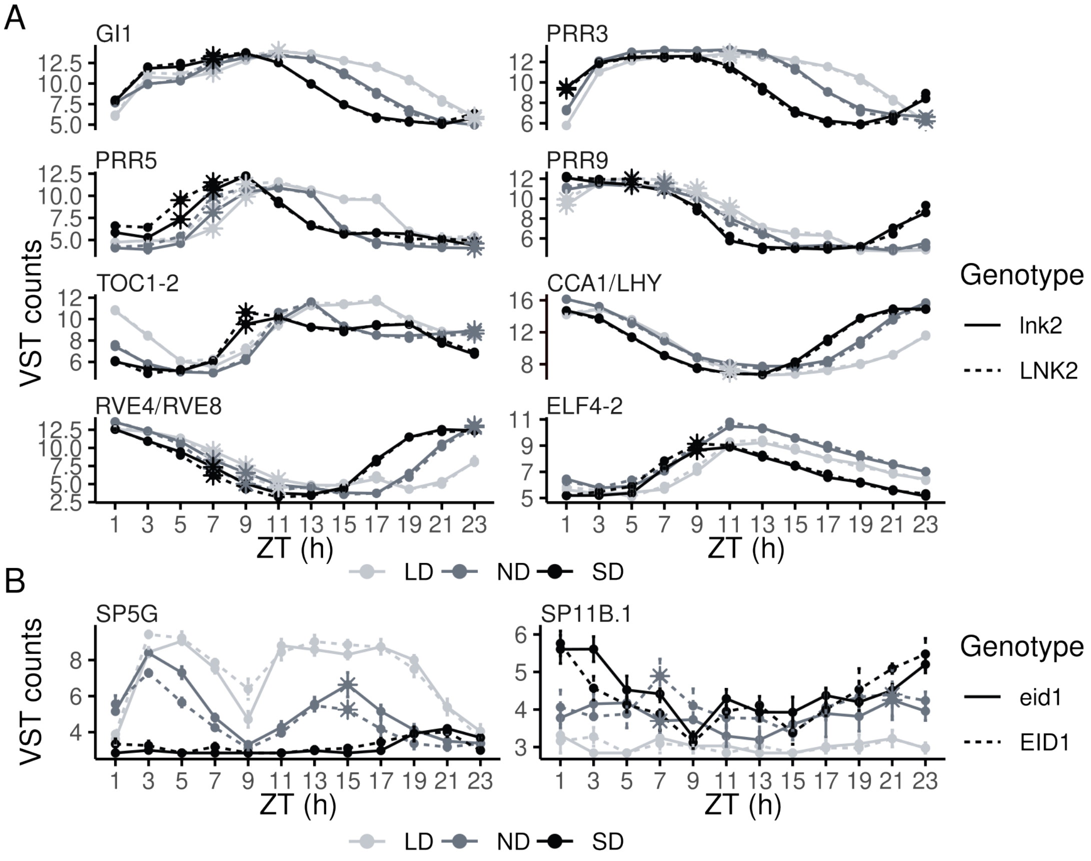


**Figure S9. Effect of LNK2 and EID1 on circadian clock and CETS transcripts.**

Average normalized expression for transcripts differentially regulated by LNK2 (A) or EID1 (B). Photoperiods are indicated by color. Asterisks mark time points with significant expression differences between genotypes grouped by allele. (A) Circadian clock transcripts differentially expressed by LNK2, and (B) CETS (CENTRORADIALIS/TERMINAL FLOWER 1/SELF-PRUNING) transcripts differentially expressed by EID1

**Supplementary Tables**

**Table S1.** List of transcripts oscillating in each photoperiod and genotype (FDR<0.01)
**Table S2.** Comparison of transcripts oscillating in our current experiment versus Müller et al., 2016.
**Table S3.** Motif Enrichment Analysis in Evening-Phased Transcripts.
**Table S4.** Motif Enrichment Analysis in Morning-Phased Transcripts.
**Table S5.** List of circadian clock and differentially expressed CENTRORADIALIS/TERMINAL FLOWER 1/SELF-PRUNING and their ITAG 4.1 annotations.
